# The Continuous Language of Protein Structure

**DOI:** 10.1101/2024.05.11.593685

**Authors:** Lukas Billera, Anton Oresten, Aron Stålmarck, Kenta Sato, Mateusz Kaduk, Ben Murrell

## Abstract

Just as language is composed of sublexical tokens that combine to form words, sentences, and paragraphs, protein backbones are composed of sub-structural elements that combine to form helices, sheets, folds, domains, and chains. Autoregressive language models operate on discrete tokens, whereas protein structure is inherently continuous, and generative approaches to protein design have borrowed more from image generation than language modeling. But autoregressive models do not inherently require their inputs and outputs to be discrete. Here we describe a generative autoregressive language model over the continuous space of protein backbones, where the distribution over the placement of each successive amino acid is conditioned on all preceding residues, and can be sampled from one residue after another. We show that this approach can learn to sample diverse and realistic protein chains, opening a new potential avenue for *in silico* protein design.

## Introduction

Generative approaches to the design of novel protein structures require placing the backbone atoms of each amino acid in three-dimensional space, such that the resulting structure is physically obtainable and energetically favorable. Current approaches include: i) ‘denoising’ models (in which we include diffusion denoising (1) and flow matching (2)), which learn to transform a noisy distribution that is easy to sample from into a target distribution that is hard to sample from; and ii) ‘deep network hallucination’ approaches (3) that use gradient descent on a pre-trained folding model. Both of these approaches are ‘holistic’ in the sense that they consider, and transform, the entire protein structure (and, sometimes, the sequence) at once.

In contrast, exceptional success has been achieved in other domains with autoregressive models that generate tokens one after another, conditioned on all preceding tokens. For text generation, ‘decoder-only’ transformer architectures (4) currently dominate. These have, at their core, a masked attention mechanism that allows each token to selectively attend to any preceding tokens when sampling the next. The masked attention structure allows the model to be trained efficiently over entire sequences in a single pass.

Such models emit ‘logits’, or unnormalized log probabilities, for each possible token, and are trained to maximize an average log probability of the training sequences. As the space over tokens is discrete, sampling from these distributions is simple: tokens can be drawn proportional to their probability, or from a ‘nucleus’ of the most probable tokens (5), which can prevent sample degeneration in an autoregressive setting. Our work began by asking whether an autoregressive approach could be used for designing the *structure* of protein backbones. While proteins fold in three dimensions, they do have an inherent linear primary ordering, where one amino acid follows another. However, while the amino acid *sequence* is discrete, and language models can be straightforwardly adapted for generating amino acid sequences (6), the placement of each amino acid in three-dimensional space is continuous. This requires two adaptations: first, the model must be able to reason about the relative placement of preceding amino acids in three-dimensional space, and second, the model must emit a continuous distribution that governs the placement of the next amino acid.

To solve the first of these problems we adapt Invariant Point Attention (IPA), which was a key innovation behind AlphaFold2 (7), to allow an autoregressive mask, ensuring that, during training, no information from a later residue can influence the placement of an earlier one. Besides this, we retain the properties of the original IPA architecture, which treats each residue as a frame (ie. with a location and rotation) and has an attention mechanism with spatially interpreted keys, queries, and values, respecting the relative geometry of the frames. When augmented with an autoregressive mask, this allows for information to propagate only from earlier to later residues in a manner that ensures invariance to rigid body transformations.

Since our model uses frames to represent residues, to solve the second problem we need to choose a representation over a set of continuous values that determines the position of the subsequent backbone frame. For protein backbones, bond lengths and bond angles are under strong physical constraints, and the primary degrees of freedom are the dihedral angles. This reduces our problem to parameterizing a sufficiently flexible (and differentiable) joint probability distribution over three continuous angles. One natural choice of a distribution over an angle is the von Mises (VM) distribution (8), which is a continuous distribution over the circle, and here we propose the Neural von Mises Mixture (NVMM) model, which maps a continuous embedding vector to a mixture of von Mises distributions over the next dihedral angle, whose log likelihood plays the same role as the log-softmax over tokens in a classical discrete transformer model.

Together, these form the basis of our Generative Invariant Angle Transformer (GIAT), which samples protein backbone structures.

## Methods

A sample is a set of residues. Each residue has:

- A frame (ie. a location and a rotation).
- A set of three dihedral angles, that determine the location of the next frame.
- A discrete amino acid character.
- A residue index number.
- A chain ID.

A GIAT (see figure 1) comprises:

**Fig. 1.**
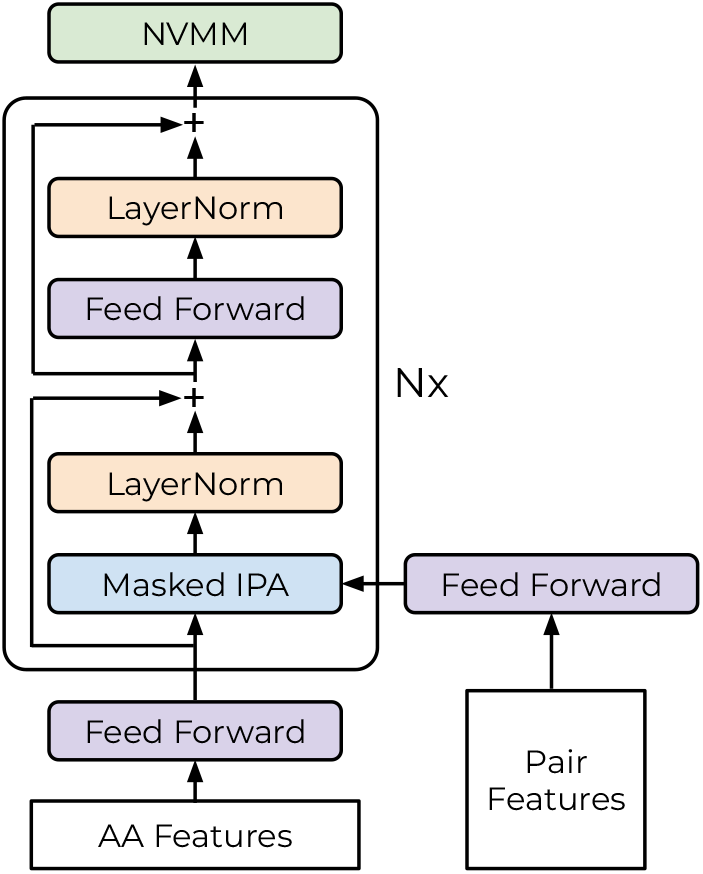
GIAT model architecture. GIAT follows a ‘decoder only’ transformer architecture, using Invariant Point Attention augmented with an autoregressive ‘causal’ mask, and using a Neural von Mises Mixture output layer that parameterizes a flexible continuous distribution over dihedral angles.

- A per-residue embedding
- A pair embedding
- A stack of masked IPA transformer layers
- A Neural von Mises Mixture layer

The output of the transformer layer is a transformed per-residue embedding, which the NVMM converts into a joint probability distribution over the dihedral angles, Ψ, Ω, Φ. Analogous to the cross-entropy loss for discrete tokens, here we use the average log probability over the observed dihedral angles. The autoregressive mask in the masked invariant point attention mechanism ensures that, during training, the location of any subsequent residues are hidden from the current residue.

### Neural von Mises Mixture

Given an embedding *S*_*r*_, the dihedral angles Ψ_*r*_, Ω_*r*_, Φ_*r*_ at residue *r* are conditionally independent of all other *S*_*t*_(*t* ≠ *r*), and all other dihedral angles (see figure 2). Thus the task of the NVMM is to place a joint distribution over *P* (Ψ_*r*_, Ω_*r*_, Φ_*r*_|*S*_*r*_). We factorize this as *P* (Φ_*r*_|*S*_*r*_, Ψ_*r*_, Ω_*r*_)*P* (Ω_*r*_|*S*_*r*_, Ψ_*r*_)*P* (Ψ_*r*_|*S*_*r*_), such that each term requires the distribution over a single angle.

**Fig. 2.**
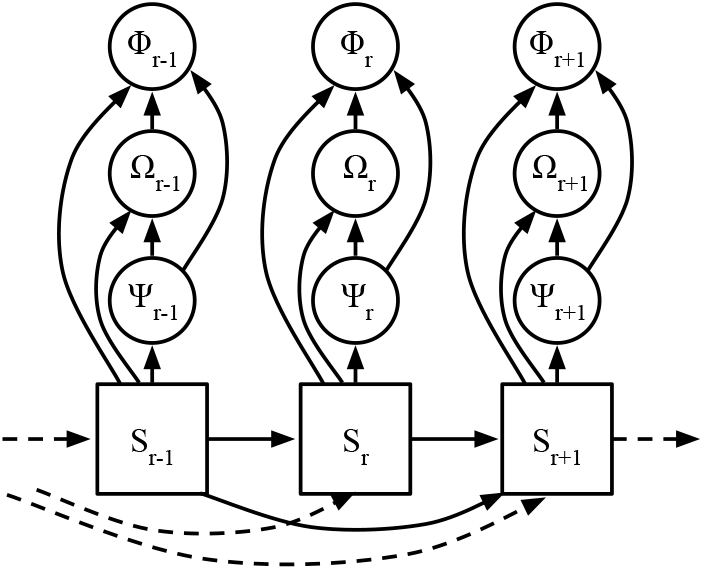
Autoregressive conditional dependence of successive residues and dihedral angles. Embeddings, *S*_*r*_ for each residue, depend upon all preceding residues. Dihedral angles, which determine the placement of the backbone of the subsequent residue, depend only upon the current residue’s embedding vector, and upon the previous dihedrals, in the order Ψ_*r*_, Ω_*r*_, Φ_*r*_

The von Mises distribution 𝒱(*μ, κ*) is a continuous unimodal probability distribution on 𝕊^1^, with density with mean *μ* and concentration *κ >* 0, where *I*_0_(*x*) is the modified Bessel function of the first kind of order zero.

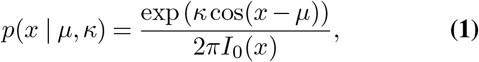

We will require out model to directly emit the VM parameters, but the standard VM parameterization does not have a smooth correspondence between the parameters and the density function. We observe, however, that the log density of the VM is a planar slice through a cylinder, and we reparameterize it such that a point on the plane controls the angular mean and concentration:

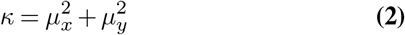

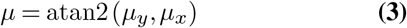

Thus when *μ*_*x*_, *μ*_*y*_ = 0 all angles are equally likely and the distribution concentrates upon atan2 (*μ*_*y*_, *μ*_*x*_) as the point *μ*_*x*_, *μ*_*y*_ moves further from the origin. Note that, for numerical reasons, we rewrite the log density to avoid atan2 (see SI section 1).

Since a unimodal distribution is insufficiently flexible for dihedral angles, we consider a weighted mixture of VM distributions, 𝒱(*μ*_x_, *μ*_y_, **w**). For a mixture of *K* components, our NVMM needs to emit *K μ*_*x*_, *K μ*_*y*_, both in (−∞, ∞), and the softmax of *K w* weight logits (see figure 3).

**Fig. 3.**
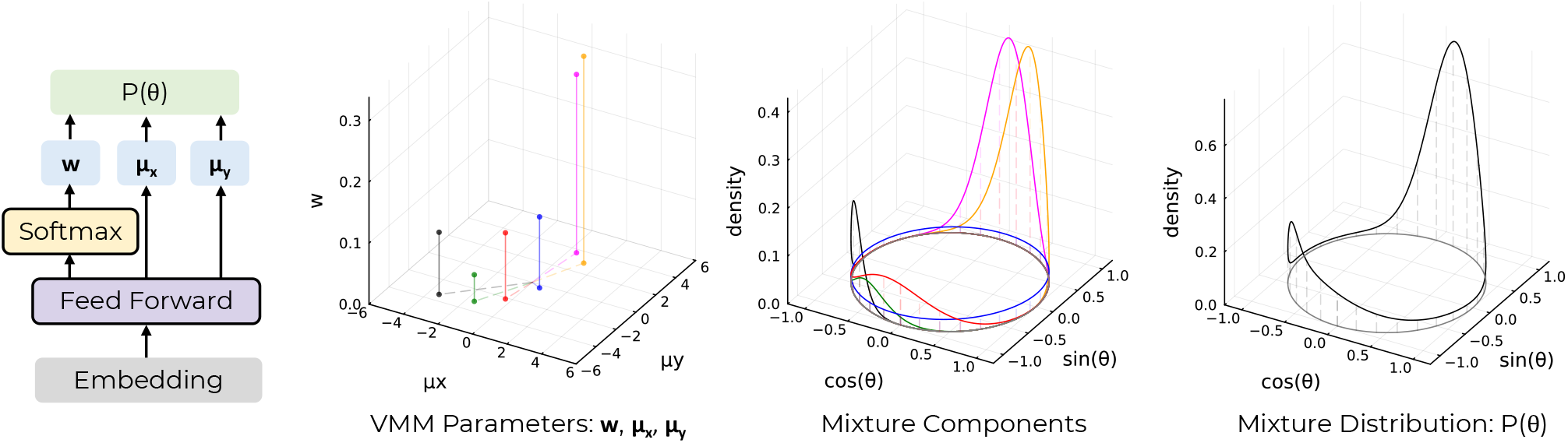
**Neural von Mises Mixture** showing (left-to-right) the architecture of the model, an example of planar VM parameters (with *K* = 6 components), the individual mixture components implied by these parameters (their colors matching), and finally the mixture density, which is the weighted sum of the components.

**Fig. 4.**
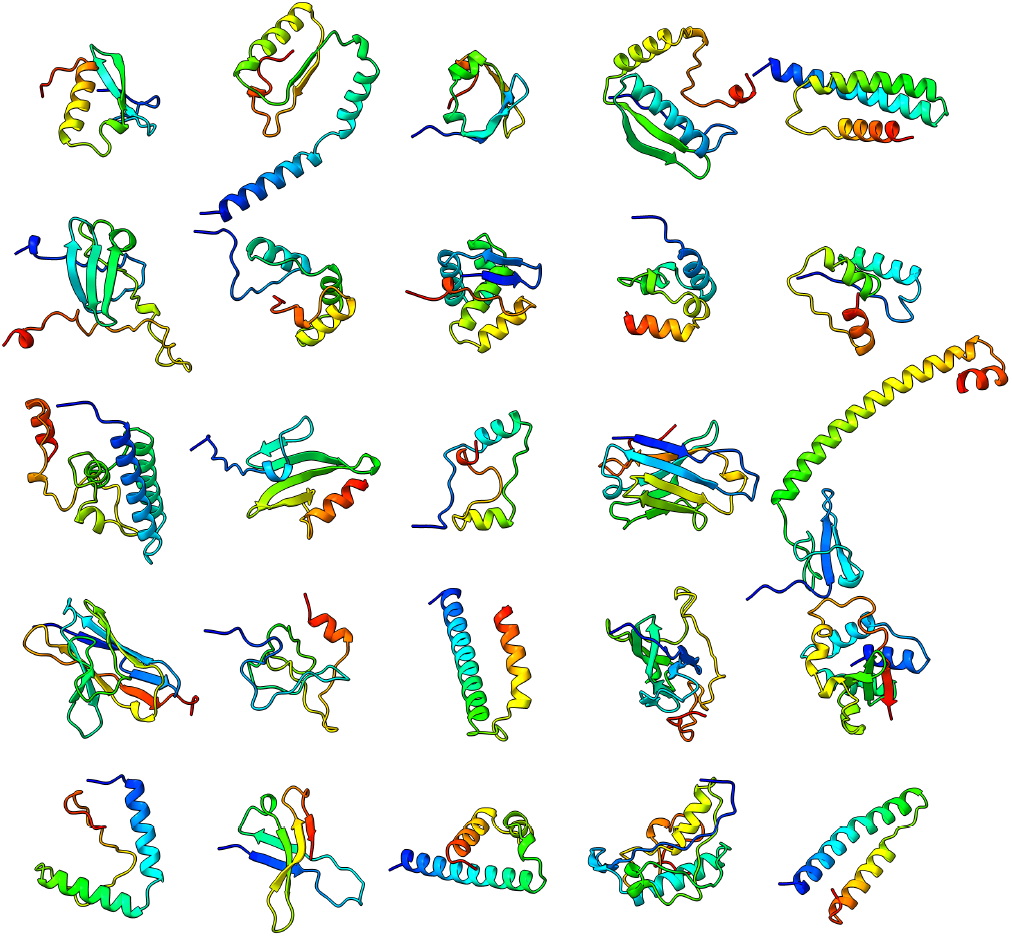
25 random samples from GIAT550, excluding pure helices, with length between 50 and 128 residues. More examples are available in figure S1.

When modeling eg. *P* (Ω_*r*_|*S*_*r*_, Ψ_*r*_), we concatenate *S*_*r*_ and *AF* (Ψ_*r*_), where *AF* is a function that converts a scalar angle to a feature vector, and proceed as above.

### Masked Invariant Point Attention

IPA, originating with AlphaFold2, augments the standard keys, queries, and values from scaled dot product attention with key points, query points, and value points, all in ℝ^3^, which are projected from the local frame of each residue. The attention weight is high when one residue’s query points are close, by Euclidean distance, to another residue’s key points. Note that, unlike AlphaFold2’s structure module, which iteratively updates the locations and rotations of the residue frames, our residue frames remain fixed during training. We introduce two modifications to IPA: 1) a causal mask, which prevents information flow from any subsequent to any preceding residue, and 2) we replace softmax with softmax1, which allows attention heads to not attend to anything (9) in a manner equivalent to having an additional ‘virtual’ zero logit. This latter change is motivated by a lack of an initial token, which can serve as an ‘attention sink’ (10) in standard large language models, but is challenging in the context of SE(3) invariance. With softmax1, the attention weights no longer sum to 1, which required modifying IPA to preserve SE(3) invariance (see SI section 2).

### Feature embeddings

IPA allows, as input, a pair representation which modulates the attention weights and informs the output embeddings. We use this to encode the difference in residue index along the primary sequence, whether or not the sequences are from the same chain (projected into a higher dimensional vector using various scalings followed by Random Fourier Features (11)), as well as to provide additional relative rotation/orientation signals, using ‘Random Orientation Features’ (see SI section 3).

Our initial residue-level embeddings, which serve as input to IPA, are a combination of a linear embedding of the amino acid one-hot encodings, added to an embedding of the relative position of each residue within a chain, which is expressed in two ways: a (scaled) number of residues until the end of the chain, and percentage of the chain that is, so far, complete. This allows the current residue to know how long the sampled chain is going to be while it is sampling successive residues. These features are noised slightly, to prevent overfitting on specific chain lengths.

### Residual Connections and Layer Normalization

Our transformer stack uses a non-standard ordering of the Layer-Norm and skip connections. We found that a standard ‘post-norm’ approach (as is used in the original GPT models (4) and in AlphaFold2) prevented scaling the number of layers in the model, whereas the standard ‘prenorm’ approach (used in more modern large language models) plateaued early during training. Instead of these, we take the LayerNorm after IPA (or Feed Forward) and sum that into the current embedding, which forces the standard deviation of that layer’s contribution to be fixed. Somewhat counterintuitively, we found this arrangement to provide favourable performance and stability over layer depth.

### Sampling amino acids

A GIAT, as depicted in figure 1 will generate a protein backbone. A sequence-given-structure model (eg. ProteinMPNN (12)) could generate an amino acid sequence conditioned on this backbone. However, applications such as ‘binder design’ might benefit from sampling a backbone conditioned on a target structure *and* sequence, since the amino acid side chains of the target might influence the binder backbone conformation. Handling this requires GIAT to jointly model sequence and structure, autoregressively. Section S3 shows how we adapt the GIAT architecture for this. The loss is then the negative mean of the dihedral log probabilities plus the cross-entropy loss of the amino acid predictions. Since the GIAT-designed sequence is only conditioned on the *preceding* backbone and sequence, we do not expect the GIAT-designed sequences to outperform state-of-the-art sequence-given-structure models, which can autoregressively design the sequence while observing the entire structure. For this reason, and in part to aid backbone benchmarking against other methods from the literature, we use ProteinMPNN for sequence generation from our backbones.

### Data preparation for training

In observed protein backbones, bond lengths and bond angles are close to idealized values, and the majority of variation in the placement of each successive backbone is determined by the three dihedral angles. We wish to model *only* dihedral angles, and ignore the variation in bond lengths and bond angles. But simply replacing the bond lengths and angles by their idealized values results in an accumulation of small errors over the length of the backbone, causing large macroscopic distortions ((13), figure S1). However, it is possible to make minute adjustments to the dihedral angles such that using idealized bond lengths and bond angles results in no accumulated distortion. This ‘idealization’ (see SI section 4) results in virtually indistinguishable structures but with only dihedral angles varying, simplifying our modeling problem.

Two data sources were used: CATH domains from AlphaFold2 models (14) and the entire protein data bank (15). PDBs from both datasets were idealized, their dihedrals calculated and their rotational frames extracted. Since residue index is used in the pair feature input, their residues were programatically renumbered by alignment to their primary sequence to remove any idiosyncrasies in the deposited PDB residue numbering. See section SI 4 for more details.

### Nucleus sampling

With autoregressive sampling, a single low probability token can derail the entire future of the sequence. Pruning regions of low probability during sampling can improve the quality of samples, and the degree of pruning can trade sample diversity against the quality of individual samples. We use a continuous variant of nucleus sampling (5). Following their notation: assuming a nucleus probability of *η* ∈ (0, 1], take *V* ^(*η*)^ ⊂ *V* = (−*π, π*) to be the smallest subset such that

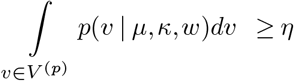

We wish to approximate sampling from:

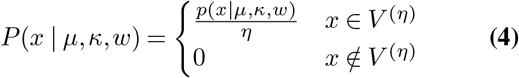

With a mixture 𝒱(*μ, κ, w*) over e.g. Θ, we first generate *N* samples *θ*_*j*_ ∼ *𝒱*(*μ, κ, w*), and compute the density at each sample:

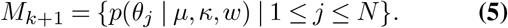

We then take Θ as the angle associated with the sample uniformly drawn from the ⌈*N · η*⌉ largest values of *M*_*k*+1_. This should approximate eq 4 as *N* → ∞.

Good choices for nucleus sampling thresholds seem to vary from one model to another, and using different thresholds for each of Ψ, Ω, and Φ sometimes appears to be beneficial. We have selected these heuristically. Interestingly, nucleus sampling can cause a kind of ‘mode collapse’ into a long pure-helix state, and for the results shown we reject samples (reporting the frequency of the exclusions) that have a contiguous alpha helix of 50 residues or longer, with no more than one non-helix residue within that window.

### Evaluation

We adopt parts of the benchmarking, evaluation, and visualization pipelines from FoldingDiff (13), using it to assess the secondary structure distributions, the diversity of samples, and assessments of re-folding using OmegaFold (16). At this point, we do not attempt extensive comparisons to other methods, and we do not expect to be state-of-the-art against Diffusion and Flow Matching models on sample generation metrics.

Following FoldingDiff, for benchmarking we sample backbones between 50 and 128 amino acids in length. Except when noted, we use nucleus sampling, and we reject pure helices (as described above), since these inflate refolding metrics as they’re easily refolded.

### Implementation

GIAT was implemented entirely in the Julia language for scientific computing (17), primarily using the Flux.jl deep learning ecosystem (18). Our implementation of Masked Invariant Point Attention is available at https://github.com/MurrellGroup/InvariantPointAttention.jl, and to accelerate autoregressive sampling, we also include a cached implementation of IPA that reduces the computation required for sampling the *N* + 1^*th*^ residue to *O*(*N*), instead of the *O*(*N* ^2^) that would be required without caching. See SI section 5. The repository with training and inference code will be available at https://github.com/MurrellGroup/CLOPS.jl. For convenience, we also provide a wrapper of the ColabMPNN Jax implementation of ProteinMPNN (https://github.com/sokrypton/ColabDesign/tree/main/mpnn (3)) so ProteinMPNN sequences can be sampled within the Julia environment: https://github.com/MurrellGroup/ColabMPNN.jl.

## Results

We explored a number of architectural variations and training strategies, and efforts to improve models are ongoing. The model whose results we describe here (GIAT550), is around 10.5 million parameters, with an embedding dimension of 196. It uses 12 IPA/feed-forward layers, with just under 80% of the parameters in the IPA layers themselves. We alternate the number of IPA attention heads, switching between 24 and 12, as well as alternating the size of the IPA ‘values’ encoding, between 8 and 16, attempting to approximately balance the per-layer memory burden. GIAT550 was pretrained, initially, on AlphaFold-predicted CATH domains, with each epoch including a single sample from each domain family, and then subsequently on the protein data bank. During training, samples are batched together to maximize GPU use, accounting for the quadratic scaling in sequence length, ie. with a larger batch dimension for shorter sequences. Structures up to 1100 amino acids were included in the training set. GIAT550 was trained on a single 48gb NVIDIA RTX 6000 Ada GPU.

GIAT550 has learnt to sample protein-like backbones with a reasonable distribution of secondary structure elements. When sequences are sampled (using ProteinMPNN) from these backbones, they are frequently predicted to re-fold into their target structure.

### GIAT learns to generate protein-like backbones

Excluding pure alpha helix samples (∼ 18% of samples, with these settings), GIAT550 appears to sample relatively compact domains. Figure 5 shows 25 random samples, with lengths between 50 and 128 residues. Further examples are shown in SI section S1.

**Fig. 5.**
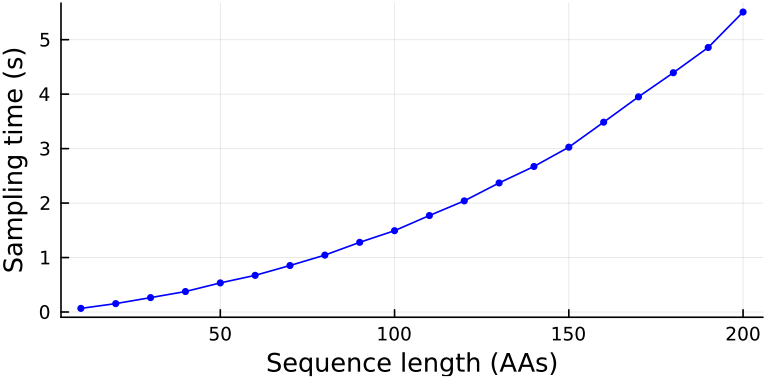
Sampling times for GIAT550, run on a CPU (Apple M3 Max), exhibiting quadratic scaling via cached IPA. Timing is the median from three runs.

The distribution of secondary structure, counting alpha helices and beta sheets, appears to approximately match that from CATH domains (see figure 6, top).

**Fig. 6.**
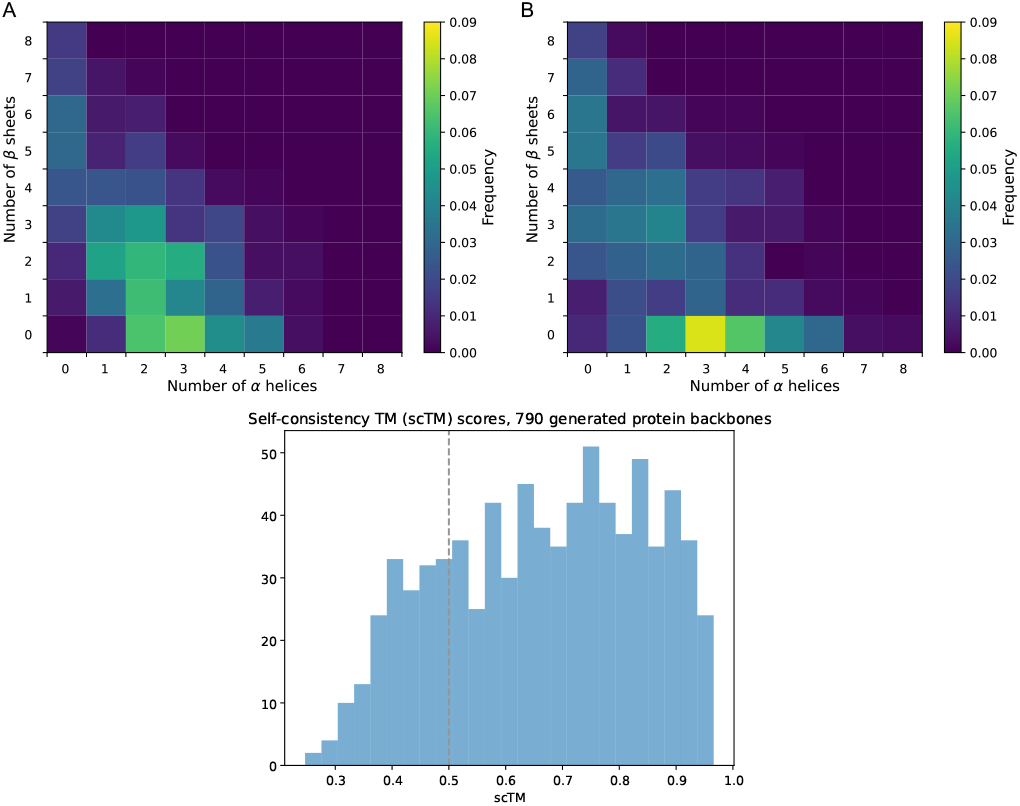
**Top, A** depicts the distribution of secondary structure elements from 780 samples taken from GIAT550 (excluding pure helices), and **Top, B** from CATH domains. **Bottom** shows the Self-consistency TM scores. Analysis and plots generated using the FoldingDiff (13) pipeline.

### GIAT models sample re-foldable backbones

Following FoldingDiff, we evaluate GIAT’s sampled backbones by *in silico* re-folding of amino acid sequences that were, themselves, sampled conditional on GIAT’s backbones. We use ProteinMPNN (12) to create eight candidate amino acid sequences which are refolded with Omegafold(16), and we take the maximum self-consistency TM-score (scTM) with the original structure across the eight candidate refolds. We perform this analysis on 780 samples from GIAT550, evenly distributed in lengths from 50 to 128, excluding pure helices. Under these conditions, the mean scTM-score is 0.65, with distribution seen in Fig 6, bottom.

### Sample quality degrades without nucleus sampling

Without nucleus sampling, we measure the decreased sample quality on three fronts, finding: 1) a higher incidence of self intersecting geometry, 2) a lower propensity for alpha helices or beta sheets, and 3) lower scTM scores (see Table 1).

**Table 1.** Comparing results from GIAT550 with unmodified sampling (ie. directly from the von Mises Mixture density), and with nucleus sampling. Using the FoldingDiff pipeline, we compare the percentage of generations without self-intersecting geometry (‘Clash-free’), the percentage of residues not contained in a helix or beta sheet (‘Loop’), and the percentage of samples where the maximum scTM-score across eight candidate sequence designs and refolds is over 0.5 (‘scTM*>*0.5’).

| GIAT550 | Clash-free | Loop | scTM>0.5 |
| --- | --- | --- | --- |
| Unmodified | 83.3% | 67% | 27% |
| Nucleus sampling | 93.2% | 45% | 78% |

### Generated backbones are diverse, but common domains are preferentially sampled

Generally, most samples from GIAT550 are distinct. Figure 7 (top) shows hierarchical clustering on structural similarity on 780 samples, with most samples appearing unrelated to one another. There are, however, some clusters. To investigate this, we identified the largest cluster, and superimposed these. Figure 7 (bottom) shows that these are immunoglobulin-like, exhibiting strong conservation in the framework regions, and high structural diversity in the complementarity determining region (CDR3). This is likely due to the fact that many protein complexes in the training dataset include antibody structures, and the model has thus learnt to sample from this common pattern a substantial fraction of the time.

**Fig. 7.**
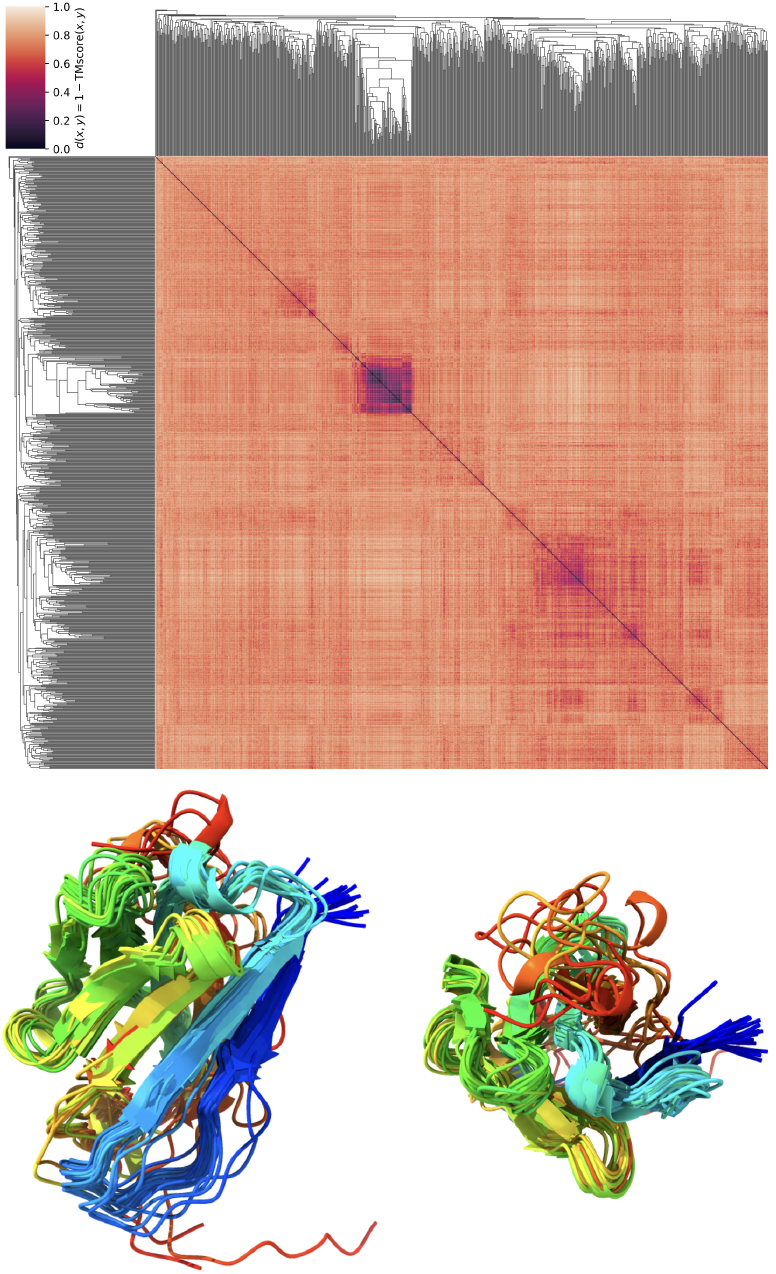
**Clustering analysis from generated samples**, (using the FoldingDiff pipeline, with distance metric *d*(*x, y*) = 1*−*TM-score(*x, y*)) showing (**top**) that, while most samples are distinct from one another, some clusters occur. **Bottom** shows structurally aligned samples from the strongest cluster, which were immunoglobulin-like, exhibiting conservation in most regions but substantial diversity in what would be their complementarity determining region (CDR) 3.

### Masked Invariant Point Attention is typically sparse, and sometimes interpretable

To understand the behavior of GIAT models, we implemented a protein backbone visualizer that can additionally overlay attention weights, depicting the degree to which the latest residue is attending to all previous residues, colored by the attention head (code at: https://github.com/MurrellGroup/ProtPlot.jl). From this, we observe that, although there are some exceptions, most attention heads are only sparsely active, attending to one or two other residues. Furthermore, most heads attend to spatially proximal residues, also with some exceptions. Sometimes, the behavior of an attention head is obvious and interpretable. For example, the top row of figure 8 shows layer 7 of GIAT550 during the design of a domain, where a single head appears to dominate, and it is active only during the construction of an anti-parallel beta sheet, only every second residue, attending only to the residue just ahead of the current one, on the opposing strand. The bottom panel shows Layer 12, which has a head that attends non-sparsely to more distant residues, possibly extracting information about the overall shape of the structure.

**Fig. 8.**
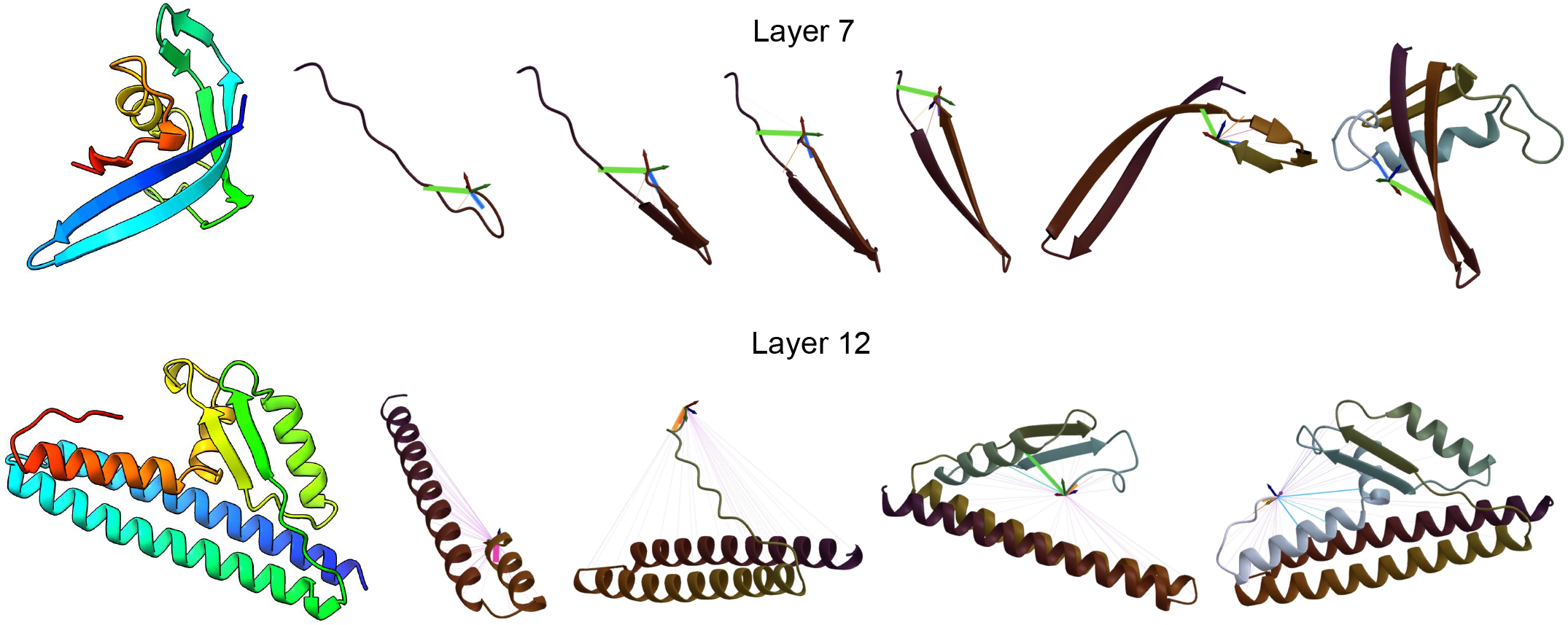
Visualization of attention head activity during design. **Top** shows a designed chain (left, rendered in ChimeraX), and a sequence of snapshots from the design process depicting Layer 7 where a single head (green line) appears to dominate, and it is active during the construction of an anti-parallel beta sheet, only every second residue, attending to the residue just ahead of the current one, on the opposing strand. **Bottom** shows an example, from Layer 12, of a more diffuse head that attends to distant residues (feint purple lines).

## Discussion

Here we have presented a new approach to generative modeling of protein backbone structures, that samples amino acid backbone placement autoregressively, one amino acid after the other. We have computationally characterized the designs from one such model, showing their diversity, and the ability of *in silico* single-sequence protein folding approaches to refold them into their target structures. We do not argue that this new approach is, at this point, going to be competitive with existing state of the art approaches for protein design, but it does suggest that this avenue of autoregressive models is worth exploring.

Interestingly, FoldingDiff (13) also describe, as a comparator for their diffusion approach, an ‘autoregressive model’ of protein backbones that generates one residue after another. However, this does not place a distribution over dihedral angles - instead it predicts the single angle that minimizes an angular loss. Without a distribution over dihedral angles there can be no stochastic sampling, and this non-probabilistic model is unable to generate any diversity. In fact it only samples the ‘most likely’ state, which is a helix. We observe this behavior in the extreme limit of nucleus sampling where we only sample from the highest mode of our angular distributions.

Given compute constraints, we have only just begun to explore the space of model architectures, with some systematic comparisons (see SI 7) justifying the modeling choices behaind GIAT550, and some more exploratory work. We have (non-systematically) explored architectures ranging from 3 IPA/feed forward layers through to 20, and parameter counts from a few million to over half a billion. Judged visually (since systematic comparisons require extensive re-folding, which competes with model development for GPU use), there appears to be substantial variation in sample quality across different models, and even within the same model from one epoch to another, often not accompanied by corresponding differences in the loss, making model selection somewhat challenging. The use of GIAT550 in particular was based on a visual impression of the samples rather than any systematic comparisons, which will follow in the future. To aid in scaling these comparisons, we have implemented custom gradients for the backward pass through IPA, which allows larger batching and faster training times than using gradients computed via Zygote.jl.

Critical future work will also explore more sophisticated sampling strategies, that consider more than just the next residue, such as continuous variants of ‘beam search’.

While some success has been had with *in silico* benchmarking of de novo designed proteins, these benchmarks are, in part, testing the ability of the folding method as well as the generative model, and protein folding from a single input sequence (rather than a multiple sequence alignment) is still unreliable. There is currently no substitute for experimental validation of designed structures, and we will be exploring this for GIAT550 and other models in this family.

Protein design, and the whole family of associated tasks, are the most obvious application for GIAT models. But GIATs are, effectively, probabilistic models over protein structure and sequence, that can be evaluated in a single pass over the entire structure, and are end-to-end differentiable. This means that they may be useful as a means to score structures generated by other approaches, and their differentiability suggests they could be used to ‘refine’ designs, by locally optimizing the dihedral angles. Another possible application for which an autoregressive model might be especially well-suited is threading an amino acid backbone through an electron density map, acting as a strong structural prior that can be rapidly sampled from, and re-weighted according to the electron density.

GIAT550 has a propensity to generate immunoglobulin domains. On the one hand this is of interest as antibody design is a key target application, and this occurring naturally in a model could be seen as promising. On the other hand, this could be considered ‘overfitting’ to some degree. To avoid models learning this sampling mode, the training data could be filtered to remove/reduce immunoglobulin chains, or to have these sampled less frequently during training, or strategies for more aggressive regularization could be explored.

The particular strengths and weaknesses of this class of autoregressive protein structure model are yet to be elucidated. Compared to diffusion and flow matching approaches, certain protein design tasks may be inherently more difficult for this kind of model, such as ‘in-painting’, where an internal region of the structure needs to be designed, conditional on a prefix and suffix backbone. It may be exceptionally difficult to learn to sample a sequence of dihedrals that result in a chain ending at precisely the right location. Another challenge, from binder design, is the placement of the *first* residue frame of the binder, which is an issue we have avoided entirely here by sampling single chains only. We are exploring strategies to overcome these issues.

Overall, these generative autoregressive transformer models over backbone structure exhibit promising initial behavior, and further exploration of model architectures, training strategies, and experimental validation will determine the place they will come to occupy in our protein design toolbox.

## ACKNOWLEDGEMENTS

This project was supported by funding from the Swedish Research Council (2018-02381 and 2023-02516) to B.M. The GPU compute for the model comparison in figure S2 was enabled by the Berzelius resource provided by the Knut and Alice Wallenberg Foundation at the National Supercomputer Centre in Sweden.

## AUTHOR CONTRIBUTIONS

Conceptualization, L.B, A.S, B.M; Formal Analysis, L.B, A.O, A.S, K.S, M.K, B.M; Investigation, L.B, A.O, A.S, K.S, M.K, B.M; Resources, B.M; Visualization, L.B, A.O, M.K, B.M; Writing – Original Draft, L.B, A.O, B.M; Writing – Review & Editing, L.B, A.O, A.S, K.S, M.K, B.M; Funding Acquisition, B.M.

## Supplementary Note 1: Numerically Stable Log PDF

In computing the log density of the von Mises distribution under our reparameterization, we avoid gradient issues associated with atan2. If *θ* = atan2(*y, x*), then *x* = *r*cos(*θ*), *y* = *r*sin(*θ*) with 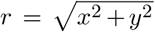. In our case, 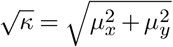 and *μ* = atan2(*μ*_*y*_, *μ*_*x*_). Applying the angle sum identity for cosine,

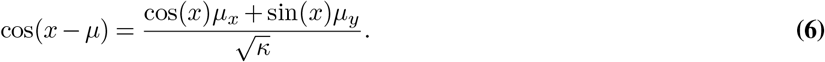

Thus,

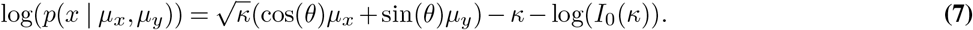

## Supplementary Note 2: Softmax1, and Invariance to Global SE(3) Transformations

We replace softmax with softmax1 in IPA, defined as

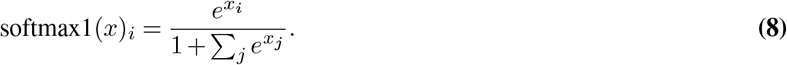

With *T*_*i*_ = (*R*_*i*_, *t*_*i*_), we modify the computation of output points to have

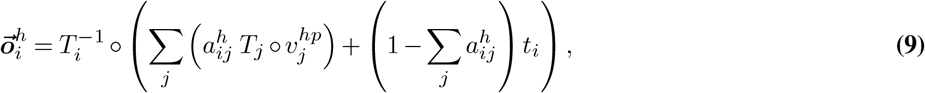

to account for the attention weights not summing to one. Now consider applying a global transformation *T*_*glob*_ = (*R, t*),

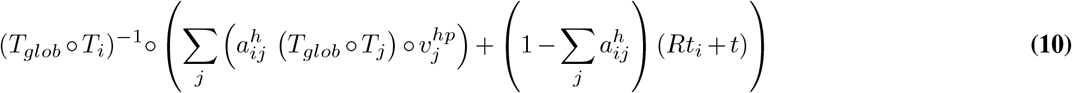

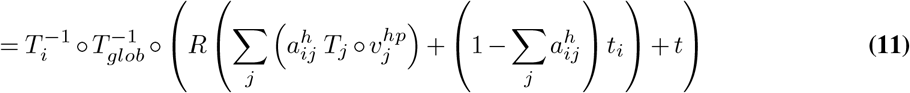

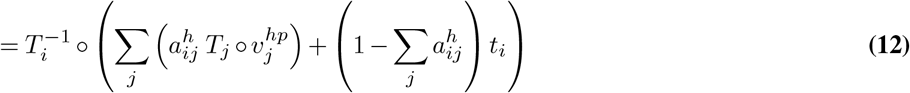

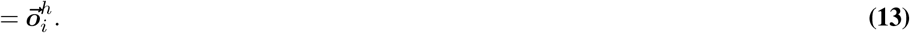

## Supplementary Note 3: Random Orientation Features

We form an initial pairwise embedding 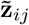 by considering distances between the projections of two sets of *d* fixed points, considered in their respective *T*_*i*_, *T*_*j*_ local frames then projected to the global frame of reference.

### Algorithm 1 Random Orientation Features

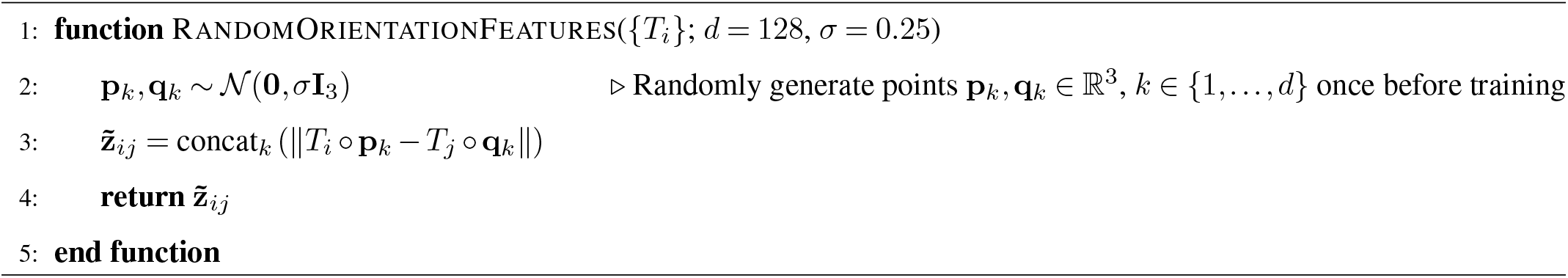

## Supplementary Note 4: Data Preparation

### A. Backbone Idealization

The goal of backbone idealization is to minimally adjust the atom positions (*N, C*_*α*_, *C*) such that the three bond lengths (*NC*_*α*_, *C*_*α*_*C, CN*) and the three bond angles (*NC*_*α*_*C, C*_*α*_*CN, CNC*_*α*_) are fixed to constant values. This allows us to represent protein backbones simply as arrays of dihedral angles.

Using a dynamic programming approach, previous work by Cui et al. (19) shows that idealized backbone structures exist with small changes to the atom coordinates. We use a gradient-based approach to the same problem, iteratively refining the position of backbone atoms under a loss function that quantifies deviations from ideal geometries, guiding the backbone towards an idealized state.

We experimented with penalizing deviations from the original atomic coordinates to prevent backbone coordinates from diverging too far, but this does not appear to be necessary and backbones efficiently converge towards an idealized nearby optimum. In cases where there are gaps in the backbone, such as when residues are missing from the structure, we mask the affected lengths and angles to prevent it from bridging the gap. Bond lengths that deviate too far (>0.5Å) from their idealized values get excluded from the loss.

### B. Residue Renumbering

Residue numbers in PDB files can be assigned arbitrarily, depending on conventions specific to particular protein families. To provide consistency to the model, we align the primary sequences obtained from the residues with a reference primary sequence, and use the new indices relative to the reference sequence as residue numbers. To ensure this alignment reflects any missing residues, we use a custom alignment algorithm with affine gap penalties that favors opening gaps in the alignment where the chain is not contiguous, i.e. when the distance between the carbonyl carbon atom of a residue and the nitrogen atom of the next is larger than expected.

## Supplementary Note 5: Cached Masked IPA

With masked IPA, we cache point and non-point keys and values during each pass-through for 𝒪(*N*) passthroughs of masked IPA. We adapt the notation used in AlphaFold2.

### Algorithm 2 Cached Masked IPA

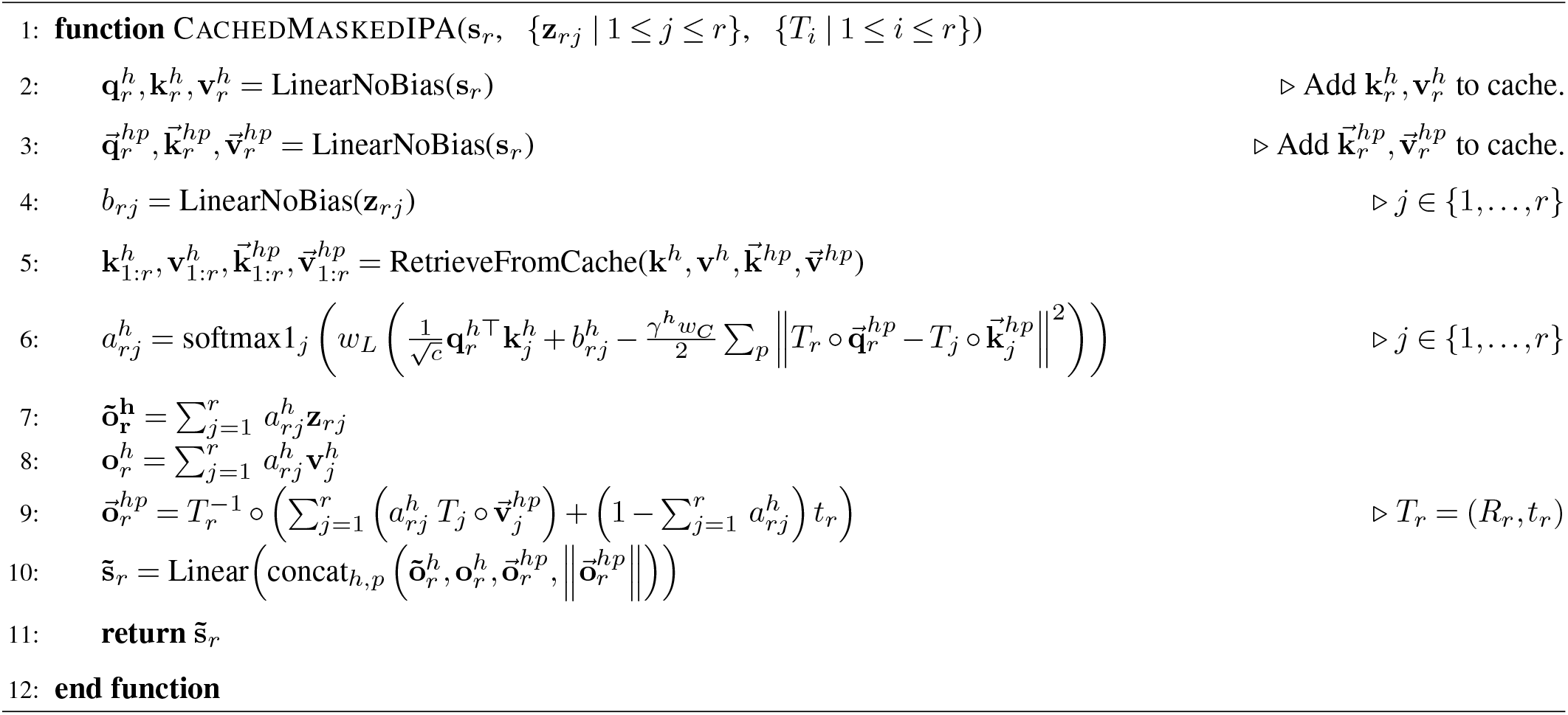

## Supplementary Note 6: Generations from GIAT550

**Fig. S1.**
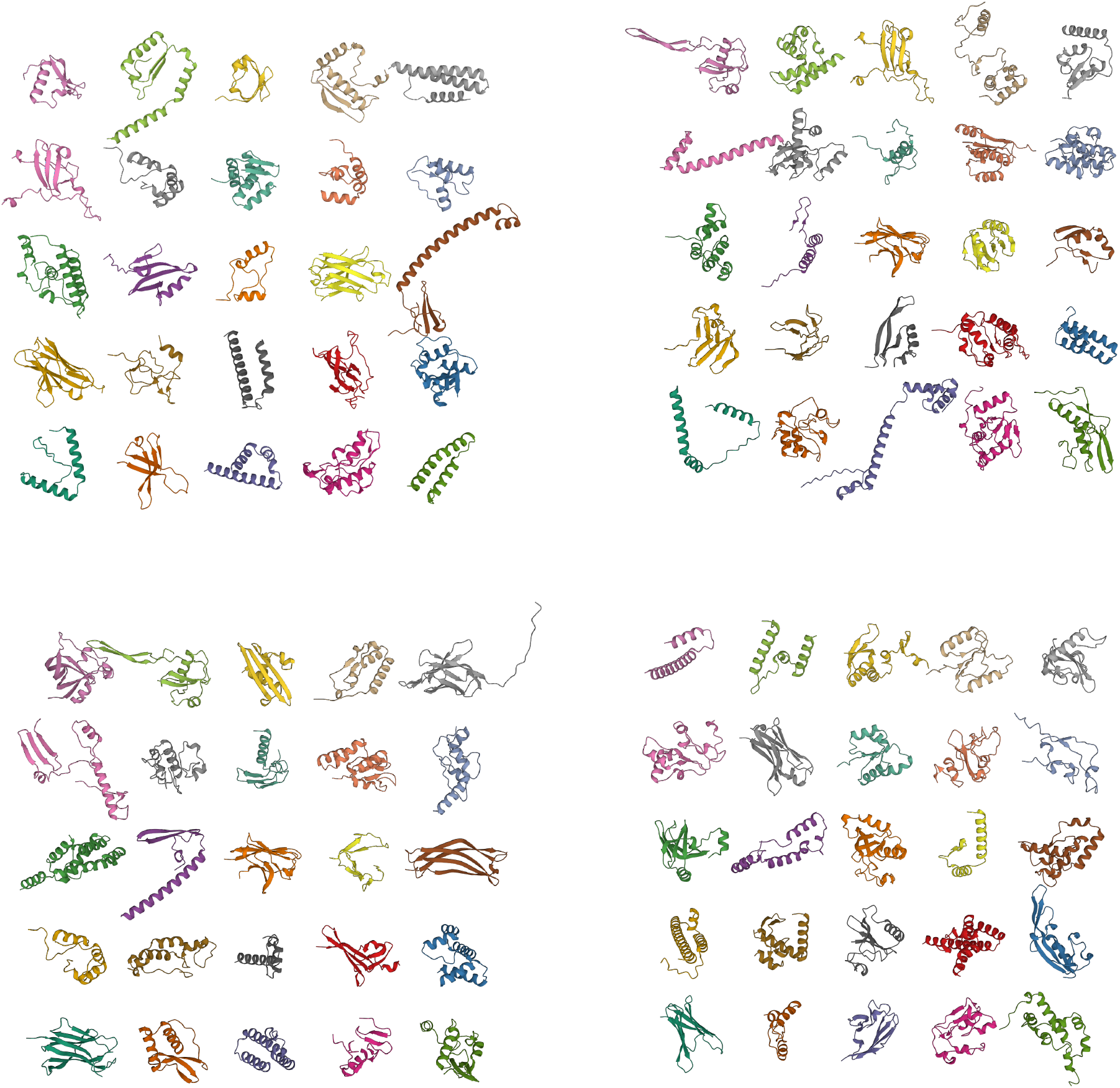
100 random samples from GIAT550, excluding pure helices, with length between 50 and 128 residues. The top left 25 samples are the same as in figure 5

## Supplementary Note 7: Model variations

Using the AlphaFold CATH domain pretraining dataset, we compared the training dynamics and loss of a number of model variations. These were run with a max chain length of 1420, on NVIDIA A100 80gb GPUs, for 500 epochs (an epoch is one sample from each domain family). Losses against one test set of holdout domain families, and one test set of PDB samples, were monitored during training.

- ‘BASE’: a model with the same architecture as GIAT550.
- ‘12LCHAINFEATS’: ‘BASE’ plus conditioning on secondary structure proportion and radius of gyration.
- ‘12LDUPLET’: ‘BASE’ but where the Φ and Ψ dihedrals are jointly sampled from a bivariate von Mises distribution, rather than sequentially.
- ‘12LMOREDROPOUT’: ‘BASE’ but with a higher dropout probability.
- ‘12LMOREJITTER’: ‘BASE’ but with larger location and rotational noise on each residue frame.
- ‘12LPOSTLN’: ‘BASE’, but where the skip connection is in the ‘post-norm’ configuration: *S* = LayerNorm(*S* + *F* (*S*)), where *F* is IPA or FeedForward.
- ‘12LSOFTMAX’: ‘BASE’ but using Softmax instead of Softmax1.
- ‘6L’: ‘BASE’ but with 6 layers instead of 12.
- ‘6LPOSTLN’: ‘6L’ with post-norm.

**Fig. S2.**
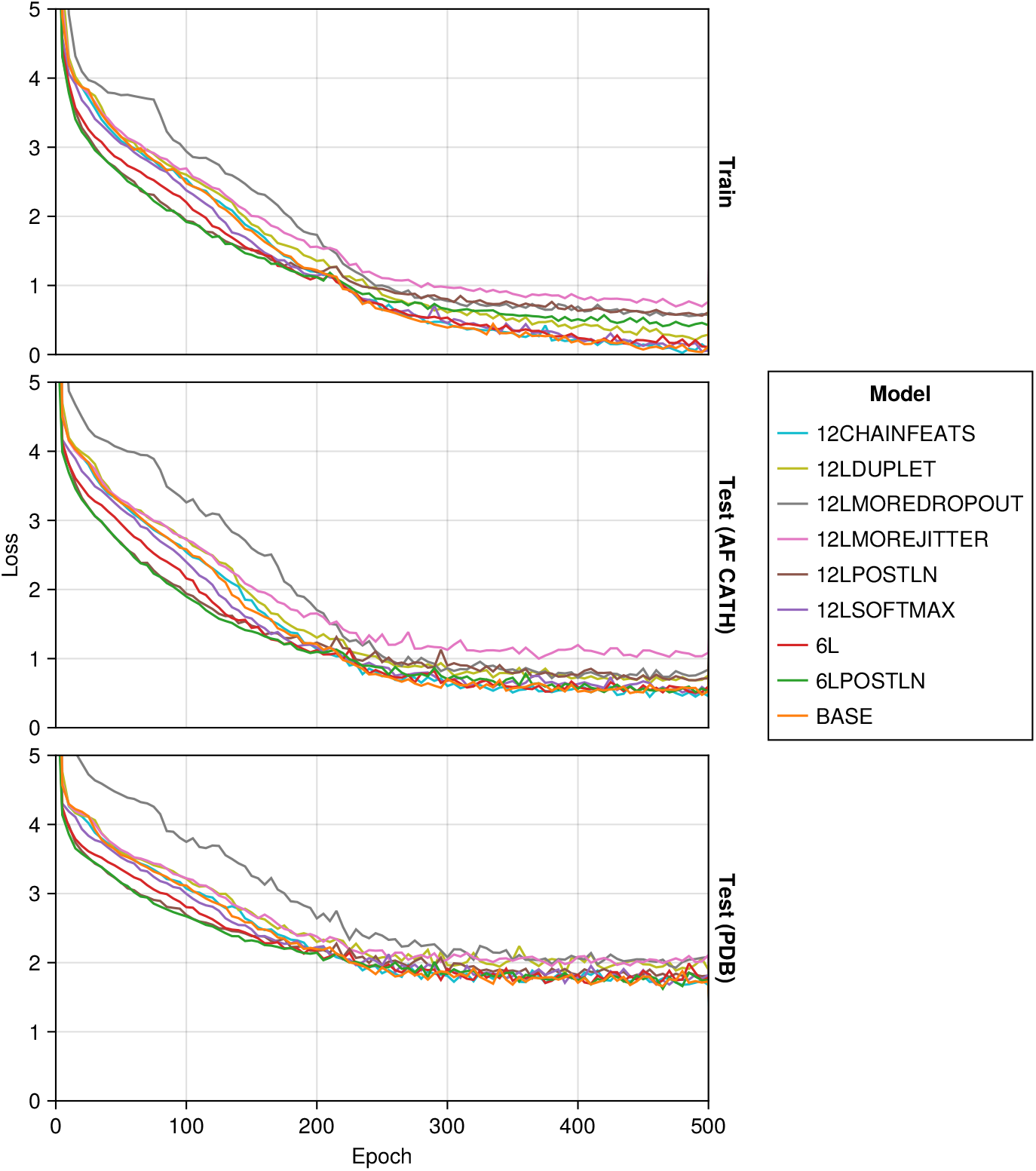
Comparing model variations.

## Supplementary Note 8: Model architecture, including amino acid sequence

**Fig. S3.**
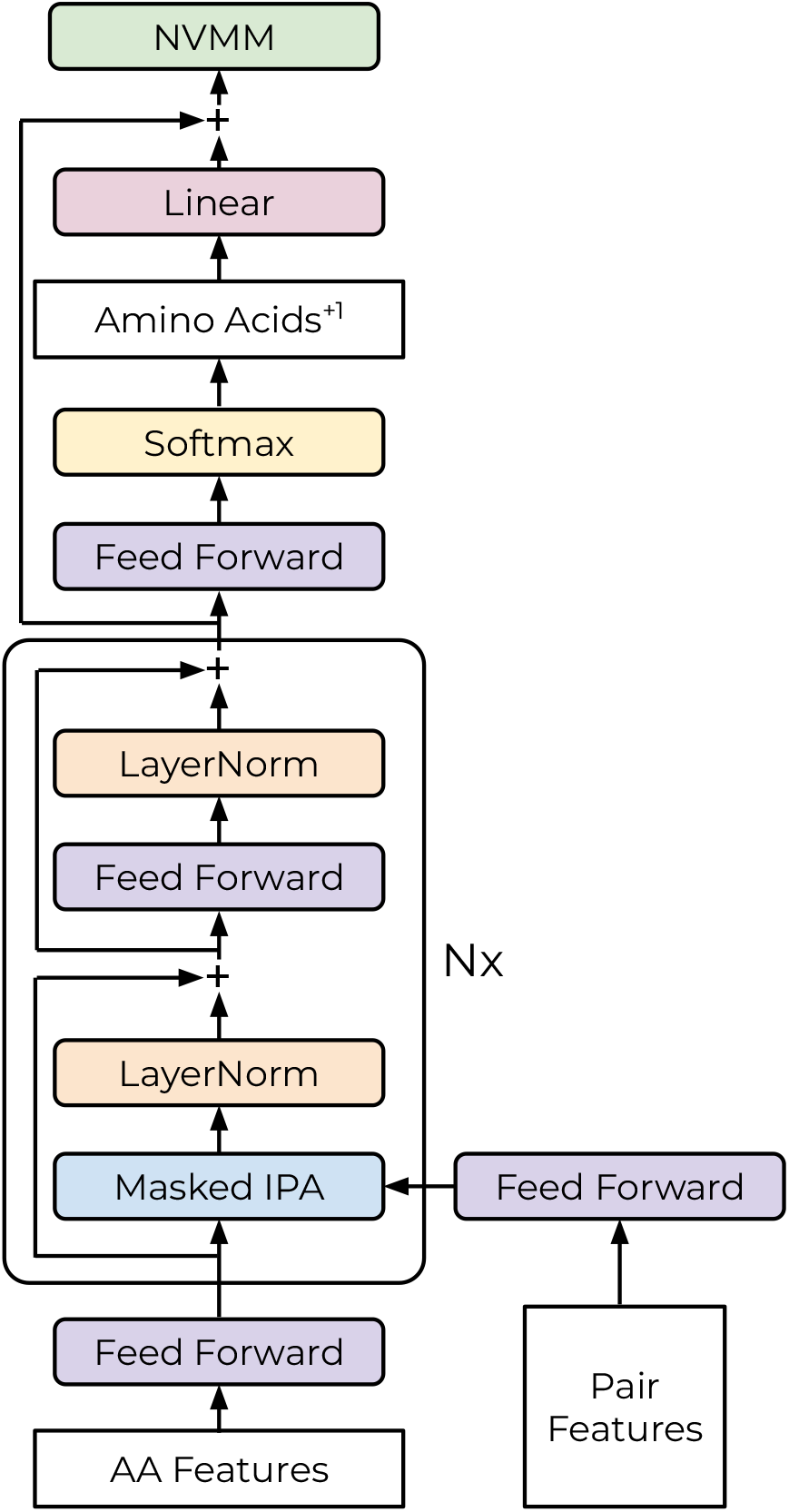
Model architecture extension to allow conditioning on, and sampling, the amino acid sequence jointly with the backbone, which is used for GIAT550. The ‘AA Features’ input also includes an embedding of the amino acid sequence, and the output from the transformer stack predicts amino acid logits of the shifted amino acid sequence (ie. over amino acid *N* + 1 at residue *N*). During training, the observed subsequent amino acid is encoded and summed into the embedding vector, which is then passed into the NVMM. During sampling, the same happens for the sampled AA. Thus the *N* ^*th*^ position samples the *N* + 1^*th*^ discrete amino acid character, and then the dihedrals that place the backbone of the *N* + 1^*th*^ amino acid.

## Bibliography

1. Joseph L Watson, David Juergens, Nathaniel R Bennett, Brian L Trippe, Jason Yim, Helen E Eisenach, Woody Ahern, Andrew J Borst, Robert J Ragotte, Lukas F Milles, et al. De novo design of protein structure and function with rfdiffusion. Nature, 620(7976):1089–1100, 2023.

2. Jason Yim, Andrew Campbell, Andrew YK Foong, Michael Gastegger, José Jiménez-Luna, Sarah Lewis, Victor Garcia Satorras, Bastiaan S Veeling, Regina Barzilay, Tommi Jaakkola, et al. Fast protein backbone generation with se (3) flow matching. arXiv preprint arXiv:2310.05297, 2023.

3. Christopher Frank, Ali Khoshouei, Yosta de Stigter, Dominik Schiewitz, Shihao Feng, Sergey Ovchinnikov, and Hendrik Dietz. Efficient and scalable de novo protein design using a relaxed sequence space. bioRxiv, pages 2023–02, 2023.

4. Alec Radford, Jeffrey Wu, Rewon Child, David Luan, Dario Amodei, Ilya Sutskever, et al. Language models are unsupervised multitask learners. OpenAI blog, 1(8):9, 2019.

5. Ari Holtzman, Jan Buys, Li Du, Maxwell Forbes, and Yejin Choi. The curious case of neural text degeneration. arXiv preprint arXiv:1904.09751, 2019.

6. Ali Madani, Ben Krause, Eric R Greene, Subu Subramanian, Benjamin P Mohr, James M Holton, Jose Luis Olmos, Caiming Xiong, Zachary Z Sun, Richard Socher, et al. Largelanguage models generate functional protein sequences across diverse families. Nature Biotechnology, 41(8):1099–1106, 2023.

7. John Jumper, Richard Evans, Alexander Pritzel, Tim Green, Michael Figurnov, Olaf Ronneberger, Kathryn Tunyasuvunakool, Russ Bates, Augustin Žídek, Anna Potapenko, et al. Highly accurate protein structure prediction with alphafold. Nature, 596(7873):583–589, 2021.

8. R von Mises. Uber die” ganzzahligkeit” der atomgewicht und verwandte fragen: Physikal, 1918.

9. Evan Miller. Attention is off by one. https://www.evanmiller.org/attention-is-off-by-one.html.

10. Guangxuan Xiao, Yuandong Tian, Beidi Chen, Song Han, and Mike Lewis. Efficient streaming language models with attention sinks. arXiv preprint arXiv:2309.17453, 2023.

11. Matthew Tancik, Pratul Srinivasan, Ben Mildenhall-, Sara Fridovich-Keil, Nithin Raghavan, Utkarsh Singhal, Ravi Ramamoorthi, Jonathan Barron, and Ren Ng. Fourier features let networks learn high frequency functions in low dimensional domains. Advances in neural information processing systems, 33:7537–7547, 2020.

12. Justas Dauparas, Ivan Anishchenko, Nathaniel Bennett, Hua Bai, Robert J Ragotte, Lukas F Milles, Basile IM Wicky, Alexis Courbet, Rob J de Haas, Neville Bethel, et al. Robust deep learning–based protein sequence design using proteinmpnn. Science, 378(6615):49–56, 2022.

13. Kevin E Wu, Kevin K Yang, Rianne van den Berg, Sarah Alamdari, James Y Zou, Alex X Lu, and Ava P Amini. Protein structure generation via folding diffusion. Nature Communications, 15(1):1059, 2024.

14. Nicola Bordin, Ian Sillitoe, Vamsi Nallapareddy, Clemens Rauer, Su Datt Lam, Vaishali P. Waman, Neeladri Sen, Micheal Heinzinger, Maria Littmann, Stephanie Kim, Sameer Velankar, Martin Steinegger, Burkhard Rost, and Christine Orengo. CATH Structural domains in AlphaFold2 models for 21 model organisms, December 2022.

15. Helen M Berman, John Westbrook, Zukang Feng, Gary Gilliland, Talapady N Bhat, Helge Weissig, Ilya N Shindyalov, and Philip E Bourne. The protein data bank. Nucleic acids research, 28(1):235–242, 2000.

16. Ruidong Wu, Fan Ding, Rui Wang, Rui Shen, Xiwen Zhang, Shitong Luo, Chenpeng Su, Zuofan Wu, Qi Xie, Bonnie Berger, et al. High-resolution de novo structure prediction from primary sequence. BioRxiv, pages 2022–07, 2022.

17. Jeff Bezanson, Stefan Karpinski, Viral B Shah, and Alan Edelman. Julia: A fast dynamic language for technical computing. arXiv preprint arXiv:1209.5145, 2012.

18. Mike Innes. Flux: Elegant machine learning with julia. Journal of Open Source Software, 2018. doi: 10.21105/joss.00602.

19. Xuefeng Cui, Shuai Cheng Li, Dongbo Bu, Babak Alipanahi, and Ming Li. Protein structure idealization: How accurately is it possible to model protein structures with dihedral angles? Algorithms for Molecular Biology, 8(1):5, Feb 2013. ISSN 1748-7188. doi: 10.1186/1748-7188-8-5.

